# Ultra-Fast Implementation of Multivariate GWAS in Genomic SEM Using Flexible Analytic Estimation

**DOI:** 10.64898/2026.06.03.729606

**Authors:** Javier de la Fuente, Mijke Rhemtulla, Travis T. Mallard, Michel Nivard, Andrew D. Grotzinger, Elliot M. Tucker-Drob

## Abstract

Many medical, physiological, and psychiatric traits and disorders are highly polygenic and exhibit complex patterns of genetic sharing and differentiation. In 2018, we introduced Genomic Structural Equation Modelling (Genomic SEM) as a formal framework and free, open source, R-based software for modelling the multivariate genetic architecture of both continuous and binary Genome-Wide Association Study (GWAS) phenotypes, interrogating their joint and distinct functional genomic pathways, and leveraging empirical models of the genetic relations among phenotypes to guide multivariate GWAS discovery. Here we introduce a closed-form analytic solution for estimating SNP effects within multivariate GWAS in Genomic SEM. This estimator is over 800 times faster than the existing iterative estimator, drastically decreasing reliance on high performance computing (HPC). On a MacBook pro laptop with M4 Max chip, a multivariate GWAS (∼1M SNPs) of 5 common factors underlying 13 phenotypes takes approximately 2 minutes. Concurrent with the release of this preprint, we are adding an analytic estimation option to the *userGWAS* function in the GenomicSEM package for alpha testing along with a tutorial on our GitHub wiki.

## INTRODUCTION

Many medical, physiological, and psychiatric phenotypes and disorders are highly polygenic and exhibit complex patterns of genetic risk sharing across a wide range of conceptually distinct diagnostic categories. With substantial investments in large consortia, registry-based efforts, and national biobanks, genome-wide association studies (GWAS) of disease traits and related quantitative phenotypes have made substantial strides in attaining the power to detect reproducible genetic associations, estimate genome-wide single nucleotide polymorphism (SNP)-based heritabilities, and estimate genetic correlations. We introduced Genomic Structural Equation Modelling (Genomic SEM)^1,2^ in 2018 to help the scientific community to more fully capitalize on this growing corpus of GWAS research output. Genomic SEM is a formal framework and free, open source, R-based software for modelling multivariate genetic architecture of systems of genetically correlated phenotypes, using such models to enhance inferential and statistical power for genome-wide discovery, and interrogating joint and distinct functional genomic pathways and external genetic correlates.

Genomic SEM enables users to model the relations among GWAS phenotypes (both binary disorders and continuous traits) based solely on publicly available GWAS summary data from participant samples of varying and unknown degrees of overlap. The reliance on GWAS summary data, which minimally consist of (linear or logistic) regression coefficients and standard errors for millions of common genetic variants, is especially conducive to general accessibility by the broad scientific community both because they are intrinsically de-identified and because they are smaller-sized files than large-scale raw genotype data (i.e., the rows of GWAS summary data correspond to genetic variants rather than individuals, such that file sizes do not increase with sample size). Currently, GWAS summary data for thousands of human complex traits can be readily downloaded from a variety of websites and repositories without prior approval. Moreover, because Genomic SEM empirically estimates the dependencies in estimation error across GWAS from the summary data, multivariate analysis using Genomic SEM is straightforward and does not require explicit knowledge of the degree to which the same participants were included in different GWAS. Because the genetic correlations and covariances that are modelled in Genomic SEM index the degree to which the true effect sizes across variants correspond across GWAS phenotypes, rather than the extent to which variation in different variables correspond across individuals, Genomic SEM can be applied to sets of variables that would be very difficult, or even impossible, to measure in the same individuals (e.g. multiple rare disorders; disorders such as bipolar disorder and major depressive disorder that are mutually exclusive of one another; early life and late life phenotypes such as childhood ADHD and Alzheimer’s disease), and sets of variables for which measurement in independent samples can aid in inference (e.g. a genetic correlation estimated using a case-control GWAS of schizophrenia and a continuous GWAS of cognitive function measured in a separate sample of healthy participants cannot be attributed to cognitive deficits induced from the disorder or its pharmacological treatment)

Some of the most inferentially powerful aspects of the Genomic SEM framework involve individually estimating the effect of each of 1M+ SNPs within a multivariate system of GWAS phenotypes (what we refer to as multivariate GWAS). However, when using the iterative estimator (from the lavaan^3^ software package) that Genomic SEM currently relies on, multivariate GWAS is computationally intensive, typically requiring access to a high-performance computing (HPC) environment. In response to this, we have now derived a closed-form analytic solution to the most computationally intensive components of the multivariate estimation process to produce dramatic gains in computational efficiency. Here we describe the analytic estimator, and report results from a series of analyses that confirm that the estimator produces near identical results (R^2^>.999) to those obtained from the original iterative estimator (we attribute the very small differences between the two estimators to numerical imprecision associated with iterative estimation). Concurrent with the release of this preprint, we are adding an analytic estimation option to the *userGWAS* function in the GenomicSEM package for alpha testing, along with an RPubs tutorial that describes the function and provides a worked example. In short, a multivariate GWAS with *userGWAS* can now be run on a standard laptop in a matter of minutes. Our intention of releasing the alpha version at this juncture is to allow interested users to work with the function and provide feedback regarding the ease of use and requested features before we advance to a more complete version suitable for beta testing.

We currently recommend that, during the alpha testing phase, users of the analytic estimator cross-validate their results against those from the iterative estimator for a small subset of SNPs (e.g. top hits and 1,000 randomly selected SNPs). The computational benefits associated with our shift from reliance on iterative estimation to a closed-form analytic solution opens exciting new opportunities for highly multivariate analyses that are currently impossible with the existing software, even when implemented within an HPC environment. However, at present, users should exercise caution when interpreting results of such analyses. We are planning to carry out validity studies for very high dimension applications in the future.

## METHODS

### Analytical Framework: Genomic SEM

Genomic SEM is a two-stage structural equation modelling approach. In Stage 1, an empirical genetic covariance matrix and its associated sampling covariance matrix are estimated by inputting GWAS summary statistics into a multivariate version of Linkage Disequilibrium Score Regression (LDSC)^4^ that accounts for varying and potentially unknown degrees of sample overlap. In Stage 2, the user specifies a structural equation model, the parameters of which are optimized to minimize a weighted fit function that indexes the discrepancy between the model-implied genetic covariance matrix and the observed genetic covariance matrix. When the effects of individual SNPs are to be included in the structural equation model (such as when seeking to perform multivariate GWAS discovery), the genetic covariance matrix estimated in Stage 1 is first expanded to include these individual SNP effects before optimization. Two key features of this multivariate discovery approach are that (a) the multivariate structure that guides the analysis is informed by the empirical genome-wide covariance structure and (b) an inferential safeguard termed the *Q*_*SNP*_ heterogeneity statistic is provided to indicate when the multivariate pattern of SNP-phenotype associations deviates from the expectations of the specified model.

#### Stage 1: LDSC model

In the first stage, we estimate the additive common variant heritability and genetic covariances using a multivariate expansion of the LDSC model^5–7^. Under LDSC, *K* phenotypes and *M* SNPs measured in *N* individuals are modelled according to the equation: *y*_*i,k*_ = *x*_*i,j*_*β*_*j,k*_ + *ϵ*_*i,k*_ where *y*_*i,k*_ is the score for person *i* on the standardized phenotype *k* (or in the case of a binary phenotype, the latent continuous liability distribution underlying propensity toward the phenotype), *x*_*i,j*_ is the standardized genotype for person *i* on SNP *j, β*_*j,k*_ is the true standardized effect size for SNP *j* on phenotype *k*, and *ϵ*_*i,k*_ is the residual for person *i* on phenotype *k*. The *β*_*j,k*_ terms are modelled as phenotype-specific random effects, varying over independent SNPs, with 𝔼[B] = 0 and *cov*[B] = (1/*M*)**S**, where **B** is the *M* × *K* matrix of true standardized genotype effect sizes for the regression of phenotype *k* on SNP *j*. **S** is a genetic covariance matrix whose diagonal elements contain the SNP-based heritabilities 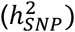 and off-diagonal elements contain the genetic covariances (*σ*_*g*_), i.e. the genetic correlations scaled relative to their SNP heritabilities.

Element *s*_*kl*_ of **S** is estimated from GWAS summary statistics by regressing the product of *Z* statistics for the linear regression of phenotypes *k* and *l* on SNP *j* on the LD score of SNP *j* and solving for *σ*_*g,kl*_ as follows:

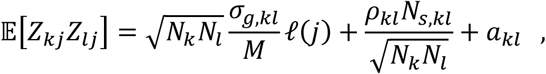

where *N*_*k*_ and *N*_*l*_ are the sample sizes for the two phenotypes, *M* is the number of SNPs, *𝓁*(*j*) is the LD score of SNP *j* (the sum of squared correlations between SNP *j* and all other SNPs in the reference panel), *N*_*s,kl*_ is the number of individuals included in both GWAS samples, *ρ*_*kl*_ is the phenotypic correlation within the overlapping participants, and *a*_*kl*_ is a term representing unmeasured sources of confounding such as shared population stratification across GWASs. Note that because the LDSC intercept is a freely estimated parameter, the *ρ, N*_*s*_, and *a* terms do not need to be known. Also note that when the *Z* statistics for the same phenotype are double entered into the left-hand side of the above equation (i.e., *k* = *l*), such that 𝔼[*Z*_*kj*_*Z*_*lj*_] becomes 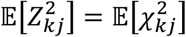, the equation reduces to the univariate LDSC model, and *σ*_*g*_ becomes an estimate of 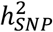.

A multivariate block jackknife, resampled across independent blocks of the genome, is used to estimate a full sampling covariance matrix, **V**, the diagonals of which contain the squared standard errors of the estimates within **S**, and the off-diagonals of which contain the sampling covariances. This jackknife has been extensively validated and found to very closely approximate the sampling covariance matrix of model parameters in raw data simulations^1,5,6^. As sample overlap creates a dependency between *Z* statistics for pairs of traits, thus increasing their products, the LDSC intercept 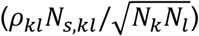 is affected, but the regression slope is unaffected such that the estimates of genetic covariance and their standard errors are unbiased irrespective of degree of sample overlap. Because the matrix of cross-trait LDSC intercepts empirically indexes the dependency in estimation errors of GWAS estimates, it serves as an estimate of the sampling correlation of GWAS estimates for the individual variants, a feature that Genomic SEM capitalizes on when individual SNP effects are later incorporated into models for multivariate discovery.

LDSC has been extensively validated and widely applied. It has been shown across a wide range of simulations to be a versatile and robust method for estimating 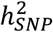, and genetic covariances *σ*_*g*_ for polygenic traits.^1,5–10^ When binary phenotypes are entered into LDSC, the SNP-heritabilities and genetic covariances are initially obtained on the observed scale, which is dependent on both sample and population prevalence, and are therefore transformed to the liability scale under a liability threshold model.^1,11^ Genomic SEM software currently includes multivariate versions of standard LDSC, annotation-stratified LDSC (S-LDSC) ^7,9^, which relaxes the assumptions of the LDSC model^12^, so as to allow for heterogeneous contributions to heritability by different gene sets and categories (i.e., functional annotations).

#### Stage 2: Estimation of the Structural Equation Model

In Stage 2, the user specifies a multivariate system of regression and covariance associations involving the genetic components of phenotypes with one another, with more general latent factors, and/or with individual SNPs. These associations are represented by a set of parameters (θ), which are estimated by minimizing a fit function indexing the discrepancy between the model-implied covariance matrix and the empirical covariance matrix obtained in the previous stage. Genomic SEM provides substantial user flexibility with respect to the particular model that is specified to produce the model-implied covariance matrix 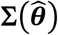 that approximates the empirical genetic covariance matrix, **S**. SEMs can be described in terms of two systems of equations, one describing the *measurement model*, and the other describing the *structural model*. In the measurement model, the genetic components of *k* indicator phenotypes are described as linear functions of a smaller set of *m* (continuous) latent variables, ***y*** = **Λη** + **ε**. In this equation, ***y*** is a *k* × 1 vector of indicators, **ε** is a *k* × 1 vector of residuals, **η** is an *m* × 1 vector of latent variables, and **Λ** is a *k* × *m* matrix of factor loadings, i.e., regressions relating the latent variables to the set of indicators. Thus, the model-implied covariance matrix of a measurement model is Σ(θ) = **ΛΨΛ**′ + **Θ**, where **Ψ** is an *m* × *m* latent variable covariance matrix and **Θ** is a *k* × *k* matrix of variances and covariances among the residuals, *ε*. To specify associations among latent variables, a structural model can be added to the measurement model to produce a full SEM. The structural model relates latent variables to each other via directed regression coefficients. It can be written in matrix notation as **η** = Γ**η** + **ζ**, where Γ is an *m* × *m* matrix of regression coefficients that relate latent variables to each other and ***ζ*** is an *m* × 1 vector of latent variable residuals. The model implied covariance matrix of observed variables is **Σ**(**θ**) = **Λ**(**I** − Γ)^−1^**Ψ**[(**I** − Γ^′^)^−1^]^′^**Λ**^′^ + **Θ**, where **I** is a *k* × *k* identity matrix.

#### Multivariate GWAS in Genomic SEM

To use the LDSC-based structural equation model to guide multivariate GWAS discovery, the ***S*** matrix is first expanded to incorporate SNP effects by appending the vector of SNP-phenotype covariances, which is directly derived from the linear or logistic (transformed to the liability scale) GWAS coefficients and the corresponding minor allele frequencies. The **V** matrix is also expanded to include the sampling covariance of the SNP effects (**V**_*SNP*_). **V**_*SNP*_ is obtained by rescaling the LDSC intercept matrix (which represents an empirical estimate of the sampling correlations of the univariate GWAS estimates) by the SEs of the SNP-phenotype covariances.

Conveniently, the sampling covariance between the LDSC-estimated genetic variances and covariances and the SNP-phenotype covariances approaches zero as the number of independent SNPs upon which LDSC is estimated increases.^1,5,6^ Thus, because of the very large number of SNPs that inform LDSC, the SNP-phenotype covariances and the LDSC-estimated genetic covariances are effectively independent. The resulting full S and V matrices are therefore:

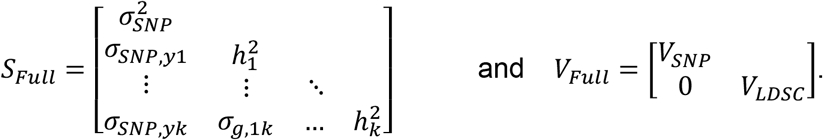

The effect of the SNP within the SEM can then be specified straightforwardly. Typically, the user specifies a *common pathways* model in which the SNP predicts each factor (**Figure 1**). This model is used to estimate effects of the SNP on each factor. Such a model is applied individually for millions of SNPs to perform multivariate GWAS. (Because it would be prohibitive to include millions of predictors in one model, and because physically distal variants are approximately independent of one another, the field standard is for GWAS— including univariate GWAS—to be run one variant at a time. The correlations among physically proximal variants, termed linkage disequilibrium, are then used to inform post-GWAS analyses, such as identification of independent loci and fine mapping).^13,14^

**Figure 1.**
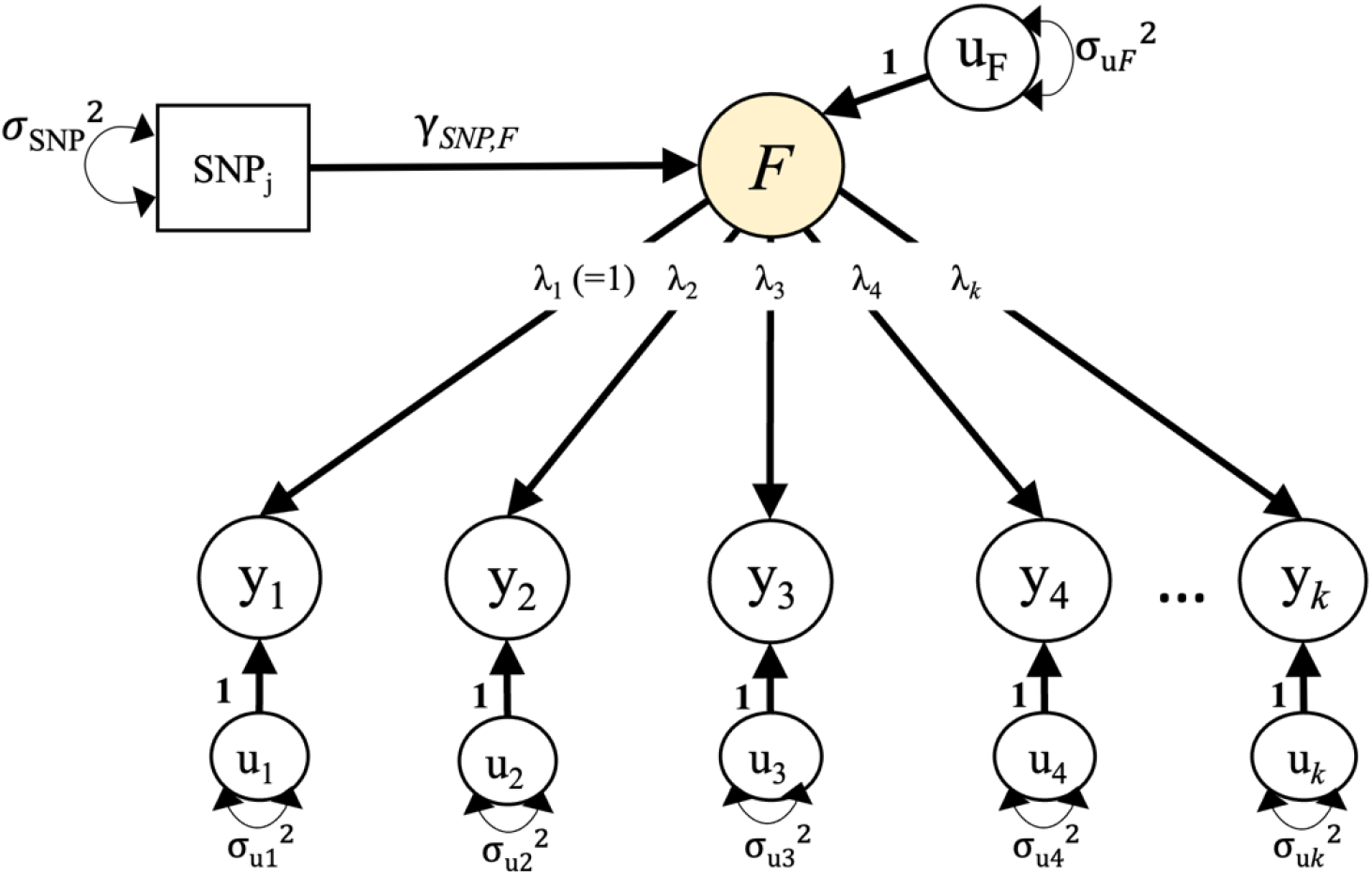
A “common pathway” model for a single common factor, with the factor F regressed on the SNP. The indicators, y_1_-y_k_, are depicted in circles, as they represent the genetic components of the GWAS phenotypes. For multivariate GWAS discovery, the model is fit individually to millions of SNPs. This model is easily extended to include multiple factors. Whereas the general matrix algebra that we use in the text employs the symbols **η, Ψ**, and **Θ** to refer to the vector of latent variables, the latent variable covariance matrix, and the covariance matrix of residuals, path diagrams often use more intuitive labels for the specific set of phenotypes and latent variables under investigation.

### Moving from Iterative Optimization to Analytic Estimation

For most SEMs, the model parameters are nonlinearly related to the observations, i.e., the elements of **Σ**(*θ*) are typically nonlinear functions of the parameter set θ. For instance, under a single common factor model, loadings *λ*_*k*_ and *λ*_*l*_ of indicators *k* and *l* on factor *η*, and the factor variance 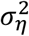, relate to the expected covariance among those indicators 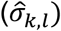 as 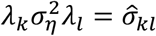. In these circumstances, a flexible general closed-form expression for the optimized parameter estimates and their SEs is not available, and an iterative estimator is used to obtain them. The general expression of the fit function that is numerically optimized is:

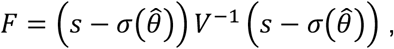

where **S** and **Σ**(*θ*) have been half-vectorized to produce *s* and 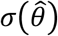, respectively. The sampling covariance matrix of the model estimates (with the squared SEs on the diagonal) are obtained as:

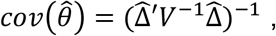

where 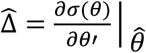 is the matrix of derivatives evaluated at the parameter estimates. To increase numerical stability given the high dimensionality of the **V** matrix, Genomic SEM employs a diagonally weighted version of the least squares fit function, in which **V** is first diagonalized to produce **V**_*d*_, and a sandwich correction is applied:

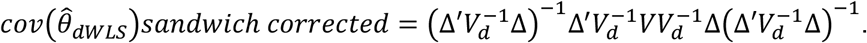

For a single model, iterative estimation is generally very fast and easily performed on a personal computer in a matter of seconds, even for a somewhat complex model. However, because multivariate GWAS requires estimation of millions individual models (one per SNP), it is prohibitive to perform multivariate GWAS using numerical optimization on a personal computer using current software.

To derive an analytic solution, we take advantage of three key observations. First, because the LDSC estimates that inform the measurement portion of the model are independent of the SNP-phenotype covariances that inform the structural portion of the model that relates the SNP to the factors^1,5,6^, they can be optimized separately. Second, whereas the measurement portion of the model requires nonlinear optimization, the structural portion of the model that relates the SNP to the factors only requires linear optimization (the observed SNP-phenotype associations, are nevertheless neither independent nor identically distributed). Third, only this structural portion of the model that requires linear optimization must be estimated separately for each SNP. The measurement model can be fixed to the estimates from the unconditional (no SNP) model such that it does not vary across SNPs, and (because of the independence of the SNP-phenotype covariances and LDSC estimates) the SEs of the SNP-factor associations remain unbiased. In fact, this helps to improve substantive inference, as it ensures that the same measurement model for the factors has been uniformly applied across all SNPs. (In practice, when measurement model parameters are freely estimated for each SNP, they are very close to those from the unconditional model).

Taken together, the computationally intensive portion of the multivariate GWAS (the portion that must be estimated separately for each of millions of SNPs) constitutes a linear system of equations, for which a flexible analytic solution is feasible. We can therefore maintain the iterative optimization for the measurement model and substitute an analytic estimator for the multivariate GWAS contained within the structural portion of the model, thereby reducing run-times dramatically. Because many different SEMs that include SNP effects (e.g. GWAS-by-subtraction, mediation models) can be reformulated as being composed of measurement models and structural models, such an approach is highly flexible and applicable to nearly all models containing SNPs for which fully iterative estimation is currently standard.

The vector of regression coefficients 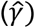 for the effects of the SNP on each factor and its sampling covariance matrix 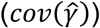, the square root of the diagonal of which contains the standard errors of the coefficients contained in 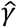) can be **estimated analytically**, using generalized least squares^15,16^ as

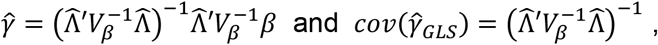

where **Λ** is the matrix of factor loadings of each indicator on each factor (with 0s entered when an indicator is not specified to load on the factor), *β* is the vector of univariate SNP-phenotype regression coefficients (directly obtained from the univariate GWAS summary statistics), and *V*_*β*_ is the sampling covariance matrix of the univariate GWAS summary statistics (obtained by transforming the LDSC intercept matrix, which represents an empirical estimate of the sampling correlation, by the SEs of the univariate SNP-phenotype regression coefficients). In practice we diagonalize *V*_*β*_ (to produce *V*_*β,d*_) to increase numerical stability. The corresponding analytic sandwich correction takes the form:

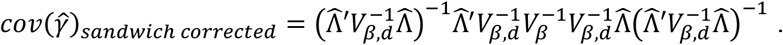

The *Q*_*SNP*_ statistic can be straightforwardly computed under this formulation. *Q*_*SNP*_ indexes the extent to which the multivariate pattern of SNP-phenotype associations deviates from the expectations of the model specified. *Q*_*SNP*_ was originally computed by comparing a common pathways model (in which SNP effects are specified to operate only on the common factors) to an independent pathways model (in which the SNP effects are specified to operate directly on the individual indicators). However, because the measurement portion of the common and independent pathways models are equivalent, they cancel out in this comparison, and we can reduce the computational burden of this calculation. *Q*_*SNP*_ is thus computed from the residuals of the structural portion of the model (i.e. the difference between observed and model-implied SNP-phenotype regression coefficients, *b* and 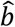), and the sampling covariance matrix (*V*_*b*_) of the SNP-phenotype regression coefficients^17^:

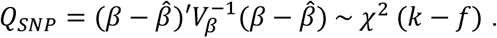

*Q*_*SNP*_ is a χ^2^ distributed test statistic with degrees of freedom equal to *k* − *f*, where *k* corresponds to the number of phenotypes and *f* corresponds to the number of factors on which a SNP effect is estimated. Factor-specific *Q*_*SNP*_ statistics can be estimated by reducing the set of SNP-phenotype regression coefficients to those corresponding to phenotypes loading only on the relevant factor.

We have added an alpha version of this analytic estimator to the *userGWAS* as an alternative to the iterative estimator provided by lavaan software.^3^ When employing the analytic option, the *userGWAS* function takes many of the same arguments as when using standard iterative estimator and continues to use the lavaan language for specifying SEMs. The lavaan language is simple and straightforward in that it does not require the end user to have knowledge of matrix algebra. For instance, similar to the way that a linear regression of an outcome on three predictors is specified in base R as ‘y ∼ x1 + x2 + x3’, a structural equation model in which one factor with three indicators is regressed on a SNP is specified in the lavaan language with the simultaneous equations ‘F =∼ y1 + y2 + y3’ and ‘F ∼ SNP’.

## RESULTS

The original 2019 paper^1^ introducing Genomic SEM and several subsequent papers^8–11,18,19^, used extensive simulation combined with empirical data analysis to validate of a wide range of properties of the iterative estimator. This included verifying that parameter estimates and standard errors are unbiased, that that the null distribution of the test statistics across replications closely match their expected distributions, that model fit statistics can be used to discern the data-generating model from plausible alternative models, and that unknown and varying amounts of sample overlap are appropriately controlled. We rely on this strong evidence base of the iterative estimator as an extensively validated benchmark against which to compare results from the analytic estimator.

As an empirical test of the equivalence of the analytic to the currently available, well-validated, iterative estimator, we drew on GWAS summary data for 13 psychiatric disorders, originally submitted to multivariate GWAS by Grotzinger et al^20^ using standard Genomic SEM software (with iterative estimation). We selected 10K SNPs at random, along with the top 2K SNPs reported for each of the 5 factors and specified the same 5-factor model specified in the original analysis (consisting of factors representing internalizing disorders, thought disorders [more recently termed the schizophrenia/bipolar factor^21^], compulsive disorders, neurodevelopmental disorders, and substance use disorders). We conducted this analysis for the selected ∼20K SNPs (SNPs total to slightly under 20K because some of the top associations overlap across factors) using the standard iterative estimator, as well as using two implementations of the analytic estimator: one that is looped across SNPs, and the other that is applied to all SNPs using vectorized code. Note that the analytic option of *userGWAS* implements the vectorized version of the analytic estimator. We conducted all analyses on a MacBook pro laptop with M4 Max chip with 16 cores and 64 GB of RAM memory.

Figure 2 illustrates the key results. It can be seen (left panel) that run times for the vectorized version of the analytic estimator were over 800 times faster relative to the standard iterative estimator, i.e. 2.5 sec vs. 2,142 sec. It can also be seen (middle panels) that multivariate GWAS beta estimates, *Z* statistics, and *Q*_*SNP*_ statistics agree nearly perfectly across iterative and analytic estimators (R^2^>.999). We attribute any minor differences that occur to arise due to approximation error in the iterative estimator.

**Figure 2.**
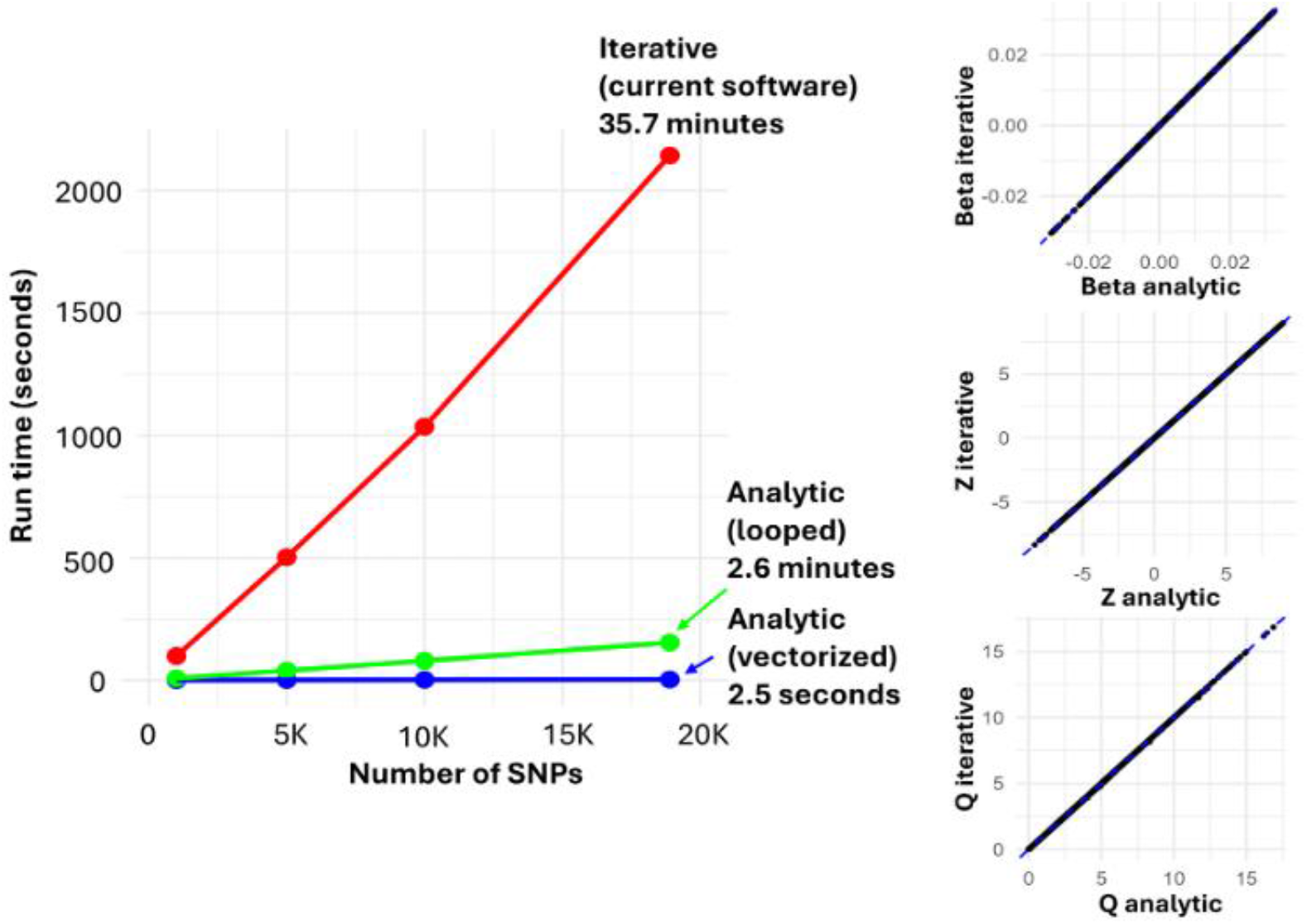
Comparison the analytic estimator and the standard iterative estimator for a 5-factor model (consisting of factors representing internalizing disorders, thought disorders [schizophrenia/bipolar], compulsive disorders, neurodevelopmental disorders, and substance use disorders) fit to 13 case-control phenotypes.^20,21^ Run time on a personal computer (Macbook Pro laptop with M4 Max chip) for the standard iterative estimator and two implementations of the analytic estimator-one that is looped across SNPs, and the other that is applied to all SNPs using vectorized code. A typical GWAS includes between ∼1M and ∼10M SNPs, such that implementation of the iterative estimator requires a computing cluster, but implementation of the analytic estimator is simple on a personal computer. On the Macbook Pro laptop with M4 Max chip, the vectorized analytic estimator took ∼2 minutes for ∼1M SNPs.

Because a typical GWAS includes ∼1M to ∼10M SNPs, the iterative estimator requires a computing cluster to estimate a full multivariate GWAS. We therefore do not compare runtimes across analytic and iterative estimators for a full multivariate GWAS, and instead focus on the runtime for a full multivariate GWAS using the vectorized analytic estimator on a MacBook pro laptop with M4 Max chip. We find that for ∼1M HapMap3 SNPs, it takes ∼2 min to run this multivariate GWAS. To avoid overtaxing available RAM, the analytic option of the *userGWAS* function runs the vectorized analysis in batches of SNPs and includes a batch size argument (current default = 100K SNPs).

## DISCUSSION

The analytic estimator now available as an option in *userGWAS* produces a dramatic reduction in the computational burden of multivariate GWAS in Genomic SEM, making genome-wide multivariate discovery feasible on standard personal computers without access to HPC infrastructure. This reduction in computational requirements does not come at the cost of inferential accuracy: empirical comparisons between the analytic and iterative estimators across ∼20K SNPs from a well-validated 5-factor model of psychiatric phenotypes produced near-perfect agreement in beta estimates, *Z* statistics, and *Q*_*SNP*_ statistics. The extremely minor differences in some estimates is likely to be attributable to numerical approximation error in the iterative estimator rather than to any deficiency of the analytic solution. Beyond democratizing access to multivariate GWAS, the shift to analytic estimation opens qualitatively new analytical possibilities: highly multivariate analyses involving hundreds of phenotypes simultaneously — currently intractable even on HPC infrastructure with iterative estimation — become computationally feasible with the analytic approach. We are actively pursuing validity studies for these high-dimensional applications.

We release the analytic option of *userGWAS* as an **alpha version** with the explicit goal of engaging the community early in its development. The function’s interface and output may change as we incorporate user feedback and extend validation work. At this stage, users should treat results as preliminary and cross-validate against the iterative option for a random subset of SNPs (e.g., ∼1,000 randomly sampled SNPs plus a sample of top hits) before drawing substantive conclusions. We strongly advise caution in high-dimensional settings involving ∼25 or more phenotypes, for which iterative cross-validation is not currently feasible. We welcome feedback on ease of use, unexpected behavior, and requested features. Users may contact the authors directly.

## Acknowledgements

EMTD is member of the Population Research Center (PRC) and the Center on Aging and Population Sciences (CAPS) at The University of Texas at Austin, which are supported by NIH grants P2CHD042849 and P30AG066614, respectively.

